# Calcium starvation leads to strain-specific gene regulation of lipid and carotenoid production in *Mucor Circinelloides*

**DOI:** 10.1101/2025.02.20.639118

**Authors:** HT Baalsrud, D Byrtusova, TH To, IE Larsen, VA Bøe, L Grønvold, J Fu, M Árnyasi, V Shapaval, SR Sandve

## Abstract

Fungi are pivotal in transitioning to a bio-based, circular economy due to their ability to transform organic material into valuable products such as organic acids, enzymes, and drugs. *Mucor circinelloides* is a model organism for studying lipogenesis and it particularly promising for its metabolic capabilities in producing oils like TAGs and carotenoids, influenced by environmental factors such as nutrient availability. Notably, strains VI04473 and FRR5020 have been identified for their potential in producing single-cell oils and carotenoids, respectively. Calcium starvation have previously been shown to have strain-specific effects, with VI04473 accumulating more lipids and FRR5020 producing more carotenoids. Here we used genome sequencing, comparative genomics, transcriptomics, and metabolite phenotyping to explore the genetic basis of lipid and carotenoid production under calcium starvation in these strains. We found extensive genomic rearrangements between these strains, as well as low conservation of gene regulatory responses to calcium depletion. This lack of conservation also applies to genes involved in lipid and carotenoid production, i.e. the lipidome. Several pathways show divergent regulation between strains, which explains the phenotypic divergence between VI04473 and FRR5020. The complex gene family evolution of several lipidome genes probably also affected specialization of these strains to calcium stress. Our study sheds light on the complexity of the evolution of metabolic networks in *M. circinelloides*. Understanding these genetic underpinnings can optimize the industrial use of *M. circinelloides*, enhancing lipid productivity and stress tolerance, and tailoring metabolic profiles for specific applications. This research advances our knowledge on complexity of fungal lipid metabolism and the role of calcium in regulating metabolic pathways.

## Introduction

Fungi play a pivotal part of transitioning from a petroleum-based to a bio-based, circular economy due to their ability to transform organic material to a variety of useful products such as food, beverages, organic acids, enzymes, antibiotics and drugs (Meyer *et al*. 2016, 2020; Dzurendova *et al*. 2022). *Mucor circinelloides* is one such species with immense industrial potential due to its diverse metabolic capabilities to produce high-value oils and other valuable compounds (Shapaval *et al*. 2014; Carvalho *et al*. 2015; Dzurendova *et al*. 2020). The versatility in metabolic pathways stems from *M. circinelloides’* adaptation to grow in a wide range of environmental conditions and substrates (Vellanki *et al*. 2018). There are numerous different strains which are ubiquitously found in soil decomposing various organic matter but can also grow on various food products as well as infect plant and animal hosts, including humans (Iturriaga *et al*. 2012; Soare *et al*. 2020; Dzurendova *et al*. 2022; Fazili *et al*. 2022). These strains vary in their ability to produce different compounds under various environmental conditions, for instance, cellular lipids can constitute between 15% (Zhang *et al*. 2007) to 54% of the total cellular weight (w/w) (Dzurendova *et al*. 2021), chitin and chitosan biopolymers can account to up to 8% (w/w) (Shapaval *et al* in prep). Although genetic tools such as CRISPR-Cas9 (Nagy *et al*. 2017, 2019) and genetic transformation (van Heeswijck and Roncero 1984; Torres-Martínez *et al*. 2012) have already been successfully demonstrated in this species, there is currently only a few strains have been sequenced, with only one annotated, long-read based genome assembly (Navarro-Mendoza *et al*. 2019). A better understanding of the genetic basis of this natural variation across multiple strains is needed to harness the biotech possibilities of *M. circinelloides* by genetically engineering strains with enhanced lipid productivity, improved stress tolerance, and tailored metabolic profiles to meet specific industrial objectives.

Two strains have recently been identified as promising industrial candidates; VI04473 for single-cell oil (SCO) production (Shapaval *et al*. 2014; Carvalho *et al*. 2015; Dzurendova *et al*. 2020) and FRR5020 for production of carotenoids (Dzurendova *et al*. 2021). SCOs are sustainable alternatives to vegetable, palm and fish oils for food and feed, and carotenoids are natural pigments with a wide range of applications (Naz *et al*. 2020). These traits can be influenced by numerous environmental factors such as type and concentration of carbon (C) and nitrogen (N) source, phosphorus (P) availability, pH and availability of metal ions such as calcium (Ca) (Gorain *et al*. 2013; Dzurendova *et al*. 2021). Nitrogen limitation conditions are known to stimulate lipid accumulation in oleaginous microorganisms where it inhibits cell proliferation and cells redirect resources from protein synthesis to lipid biosynthesis (Carvalho *et al*. 2015). Interestingly, Ca starvation was shown to have strain-specific effects on lipid and carotenoid accumulation; in VI04473 Ca starvation led to a higher accumulation of lipids, whereas in FRR5020 there was small changes in lipid content but a large increase in carotenoid content (Dzurendova *et al*. 2021). These effects were also partially dependent on phosphorous concentration and thus pH, which is known to affect enzyme activity, nutrient availability, and metabolic pathways (Bhalla *et al*. 2022). The enhanced production of lipids and carotenoids was triggered at low levels of phosphate. This is of particular interest to industry, as phosphate is a limited resource (Drangert 2012) and optimizing the production of valuable metabolites under nutrient poor conditions is hugely advantageous. A significant knowledge gap towards this end is understanding the genetic basis of metabolite accumulation in diverse *M. circinelloides* strains.

*M. circinelloides* is an emerging model for understanding lipid metabolism and a catalogue of ∼200 genes that are involved in lipid synthesis – hereafter denoted as the lipidome – have been described for various strains (Tang *et al*. 2015a; b; Sokołowska *et al*. 2023). These studies have revealed that many of the gene families in the lipidome have complex relationships characterized by gene/genome duplications and losses, which could greatly contribute to phenotypic differences between strains (Sokołowska *et al*. 2023). Differences between strains could also stem from utilizing different metabolic pathways. For instance, there is a trade-off between producing lipids and carotenoids, as both the mevalonate pathway (lipid and carotenoid) and TAG pathway (lipids) use the same precursor, Acetyl-CoA. These pathways and lipidome genes in general are well-described in various fungi (Lu *et al*. 2020; Yook *et al*. 2025). However, we lack understanding of the genetic basis for strain differences and how these differences are related to their divergent response to Ca starvation.

Ca starvation led to increased lipid accumulation in highly diverged species such as oleaginous algae (Gorain *et al*. 2013), mammalian adipocyte cells (Wang *et al*. 2017), and Mucoromycota fungi (Dzurendova *et al*. 2021). However, the genes involved in Ca signaling are not the same in these different systems; in mammals the Ca-sensing receptor plays a crucial role whereas in fungi the Ca/calcineurin pathway is the main respondent to extra and intracellular Ca levels. Calcineurin also affects the switch between different growth forms in *M. circinelloides*, from filamentous hyphae, yeast-like structures, to multinucleate sporangia (Lee *et al*. 2013). Ca could thus have wide-ranging and complex effects on growth and metabolism. Knowing that the FRR5020 and VI 04473 strains phenotypically respond quite differently to Ca availability a detailed study of their genetic and gene regulatory differences could shed light on the role of Ca in lipid metabolism in *M. circinelloides*.

In this study we combined genome sequencing, comparative genomics, transcriptomics and metabolite phenotyping to investigate the molecular basis for lipid and carotenoid production in two phenotypically divergent strains of *M. circinelloides*; FRR5020 and VI04473. We find large differences in the genome structure of *M. circinelloides* strains, as well as potentially causal differences in gene content and gene regulation underlying the extreme strain variability in metabolite production.

## Results

### Phenotypic divergence between strains in response to Ca starvation

To characterize lipid and carotenoid production in VI04473 and FRR5020 we applied the same methods as previously described by (Dzurendova *et al*. 2021). The average lipid content ranged from 24-59% of the dry cell weight (dcw), with the highest lipid content found in VI04473 (Figure 2 A and D). As expected, we observed marked differences in lipid accumulation ability between strains. VI04473 displayed Ca-dependent changes in lipid accumulation with negligible effects of different P levels. At a Ca starvation (Ca0) and P limitation (P0.5), VI04473 displayed an increase in lipid content from 34 to 51 % of DCW, while at other P levels the increase in lipid content was not observed as it was reported by (Dzurendova *et al*. 2021). Lipid accumulation in FRR5020 on the other hand is not influenced by Ca availability but decreases at high levels of P (Figure 2D). Biomass production was in accordance with the previously published by (Dzurendova *et al*. 2021), where decrease in P resulted in the decrease in biomass for both strains when Ca was present. The same was observed FRR5020 under Ca deficiency, while it was relatively stable for VI04473. The lipids-to-proteins ratio was also affected positively by Ca starvation at P0.5 in VI04473, with no effect at P1 and P4 (Figure 1B). In FRR5020 there was large variability in lipids-to-proteins ratio, with no apparent effect of Ca (Figure 1E). The level of Ca did not have any discernible effect on the fatty acid profiles (Supporting information). In summary, our experiments reproduced previous findings (Dzurendova *et al*. 2021) of strong stain-specific differences in Ca-starvation dependent lipid accumulation, but only at low levels of P.

**Figure 1:**
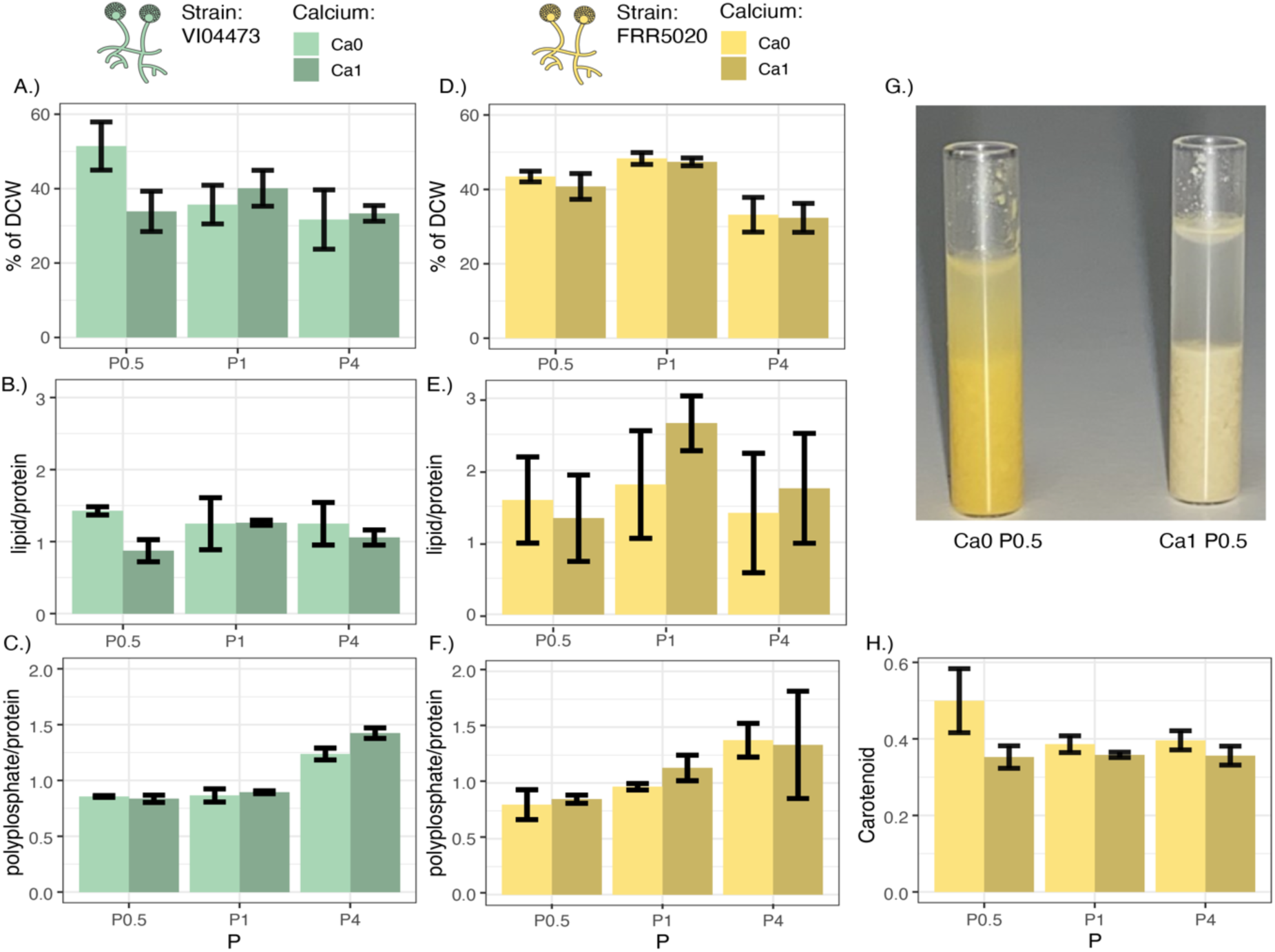
The effect of Ca and P on lipids, proteins, polyphosphates and carotenoids in *Mucor circinelloides*. Strain (FRR5020 and VI04473) and Ca (Ca) level is colored according to the legend, with Ca1 being the reference level of Ca and Ca0 is absence of Ca in the media. Phophate (P) has three levels, P1 is the reference level and P0.5 and P4 is half or four times that level, respectively. A.) and D.) shows lipid content as a percentage of dry cellular weight (DCW), B.) and E.) is the lipids-to-protein ratio, C.) and F.) is the polyphosphates-to-proteins ratio, G.) is a picture of tubes containing tissue of FRR5020 with or without Ca at P0.5 level and H.) is the level of carotenoids in the FRR5020 strain quantified by the RI_1523_/RI_1445_ ratio. For pictures of all replicates see Supporting information.

**Figure 2:**
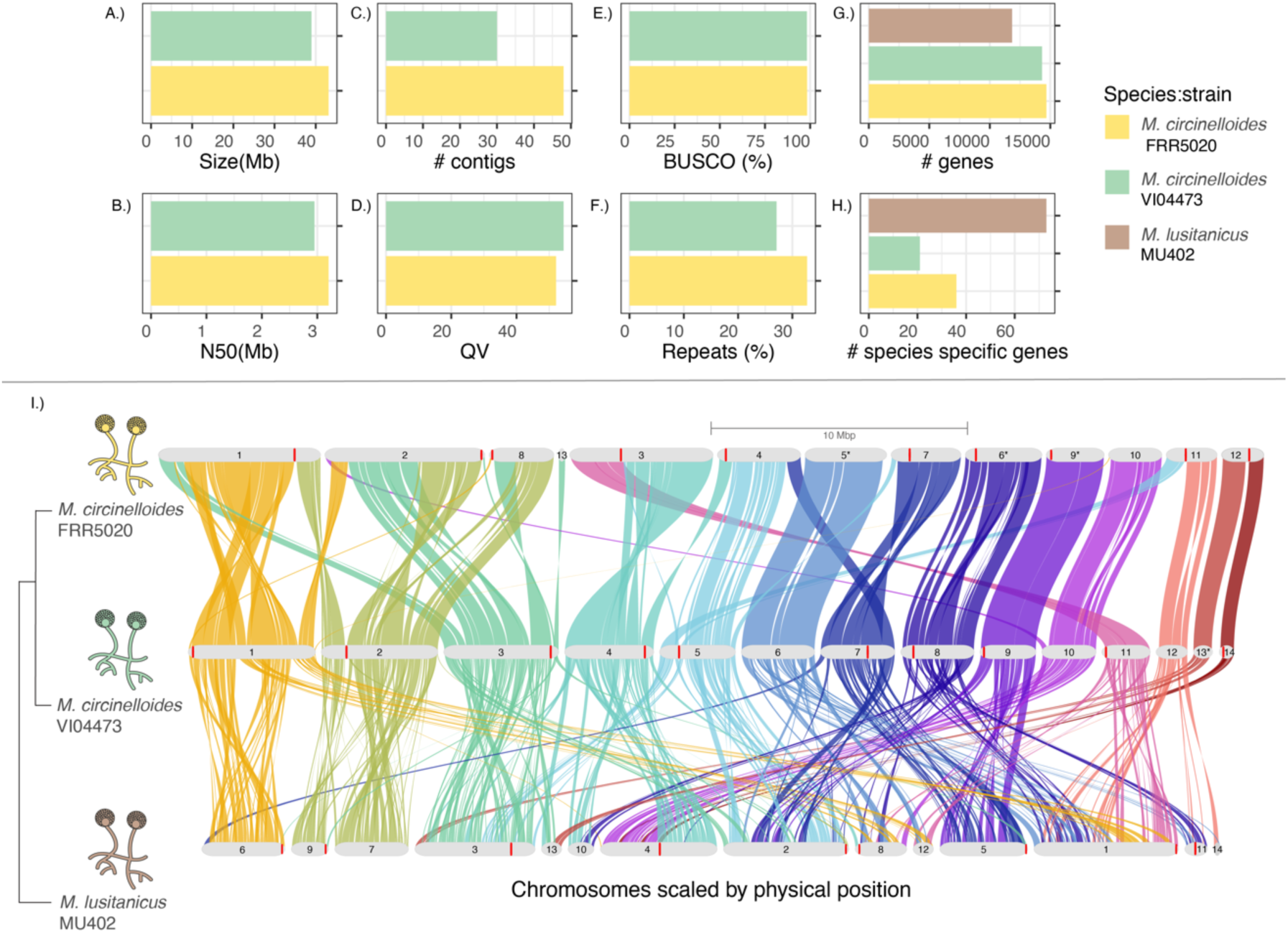
Mucor genomes. Barplots showing assembly statistics for the two *M. circinelloides* strains, FRR5020 and VI04473, including A.) genome size (Mb), B.) contig N50 (MB), C.) total number of contigs, D.) assembly quality value (QV), E.) BUSCO completeness score and F.) repeat content. Comparative genomic statistics for FRR5020 and VI04473, and Mucor *lusitanicus* strain MU402 with G.) the total number of genes for each strain, H.) the number of species-specific genes for each strain, and I.) Genome-wide synteny relationships between the three strains. The plot was generated by GENESPACE **(Lovell *et al*. 2022)**. The phylogenetic tree was generated by Orthofinder (see Supporting information for the full tree). Chromosomes are ordered horizontally to maximize collinearity with the genome assembly for the VI04473 strain. Only chromosomes or scaffolds with >100 genes and collinearity blocks > 5 genes were included in the plot. The size of the putative chromosomes are scaled by Mbs according to legend. Braids illustrate gene order along the chromosome sequence. *Indicate inverted chromosomes.

For the high pigment producing strain, FRR5020, there was a significant effect of Ca starvation on pigmentation levels (Figure 1G-H). Carotenoid content in Ca0P0.5 was significantly higher than for all other treatments, with a mean value of RI_1523/RI_1445 of 0,5 (Figure 1G). This effect of Ca at the P0.5 level is highly visible, with biomass grown at Ca0 being a bright yellow color as opposed to milky yellow at Ca1 (Figure 1H).

### Chromosome-level assemblies for two *Mucor circinelloides* strains

To investigate genetic differences between strains we generated highly contiguous genome assemblies meeting the quality standards of the Earth Biogenome Project for both VI04473 and FRR5020, with contig N50 ≥ 3Mb, QV ≥ 47 and BUSCO ≥ 98 (Supporting information). Canu generated better assemblies than Flye, possibly because of the step in Canu that performs error-correction of reads. Total length of the two assemblies were very similar, 43 and 39 Mb for FRR5020 and VI04473, respectively, which is slightly larger than other high-quality genome of *M. lusitanicus* strain MU402 (37 Mb) (Navarro-Mendoza *et al*. 2019). The Canu assemblies were annotated using RNA seq data in the funannotate pipeline, resulting in 14677 genes in FRR5020 and 14313 genes in VI04473. Karyotype is unknown in this species, but the highly contiguous assemblies contained 13 (FRR5020) and 14 (VI04473) gene-containing contigs. We also annotated centromeres based on homology with centromere sequence motifs from MU402 (Navarro-Mendoza *et al*. 2019) (Figure 2).

### Extensive genomic rearrangements between *Mucor circinelloides* strains

We used Orthofinder (Emms and Kelly 2019) to determine the phylogenetic relationship between the VI04473 and FRR5020 strains and compare gene family content using all protein-coding genes across all the publicly available annotated Mucoromycota genomes as well as outgroups (30 species/strains in total, see supporting information). Phylogenomic analyses shows that FRR5020 is the sister to *M. circinelloides* 1006phl, and both form a monophyletic clade with VI04473 (Fig. S2). *M. lusitanicus* and *M. ambiguus* form the sister group to *M. circinelloides*.

The gene content is highly similar between VI04473 and FRR5020. Given the collection of 30 fungi genomes available, we identified 13 species-specific orthogroups containing 36 species-specific genes in The FRR5020 genome, and 10 species-specific orthogroups containing 21 species-specific genes in the VI04473 genome. Figure 2 and Supporting information).

We also investigated genome-wide synteny between the two *M. circinelloides* strains and one *M. lusitanicus* strain with high quality genome assemblies (FRR5020, VI04473 and MU402) (Fig. X). A large amount of rearrangement could be seen between MU402 and our *M. circinelloides* assemblies while the synteny between FRR5020 and VI04473 was much more conserved.

### Low conservation of gene expression response to Ca starvation between strains

We carried out differential expression analyses for each strain separately to identify which genes that are differently expressed in response to Ca starvation. As the main phenotypic differences in lipid and carotenoid accumulation between strains are between growth media with Ca0P0.5 and Ca1P0.5 (Figure 1), we focused on this contrast to investigate the effect of Ca starvation on gene expression. This also allowed us to combine expression data from two independent experiments (Supporting information). For each strain we performed differential expression analysis using DESeq2 (Love *et al*. 2014) to identify differentially expressed genes (DEGs) with a model including Ca level and experiment as factors. For VI04473 we obtained 11 653 genes with sufficient RNA-seq data, with 1 524 DEGs being upregulated and 1 743 DEGs being downregulated in response to Ca starvation (padj < 0.1)(Supporting information). For FRR5020 we obtained 11 765 genes with sufficient RNA-seq data (DEGs; padj < 0.1), with 1 752 DEGs being upregulated and 2 016 DEGs being downregulated (padj < 0.1)(Supporting information).

To identify gene regulatory differences that could explain strain differences in the phenotypic response to Ca starvation we first compared log2 fold changes (LFC) in gene expression for all 1:1 orthologous genes. There were 10 483 1:1 orthologs with sufficient RNA-seq data across strains (For overall overview of ortholog relationships see Supporting information). Surprisingly, there was no correlation between LFC for the two strains (slope = 0.047, p = 2.382x10^-11^, R^2^ = 0.004) (Figure 3). The overall majority of the statistically significant DEGs (padj < 0.1) were only significant in only one strain (4 920 out of 5 999). 437 of the DEGs were significant in both strains but displayed opposite direction of LFC. Only 642 significant DEGs show the same direction of expression change in the two strains, with 232 being downregulated and 437 being upregulated under Ca starvation (Figure 3). These results show that the genome-wide gene regulatory response to Ca starvation is highly strain-specific with only a minority of genes with conserved direction of gene expression. The lack of conservation could be because Ca starvation is particularly stressful for the cells triggering a chaotic gene expression response. To test this hypothesis we also checked the between-strains correlation of gene expression between the P1 and P4 treatments, which both represent less stressful conditions for fungi. We found a significant positive correlation of 0.63 between expression (Supporting information).

**Figure 3:**
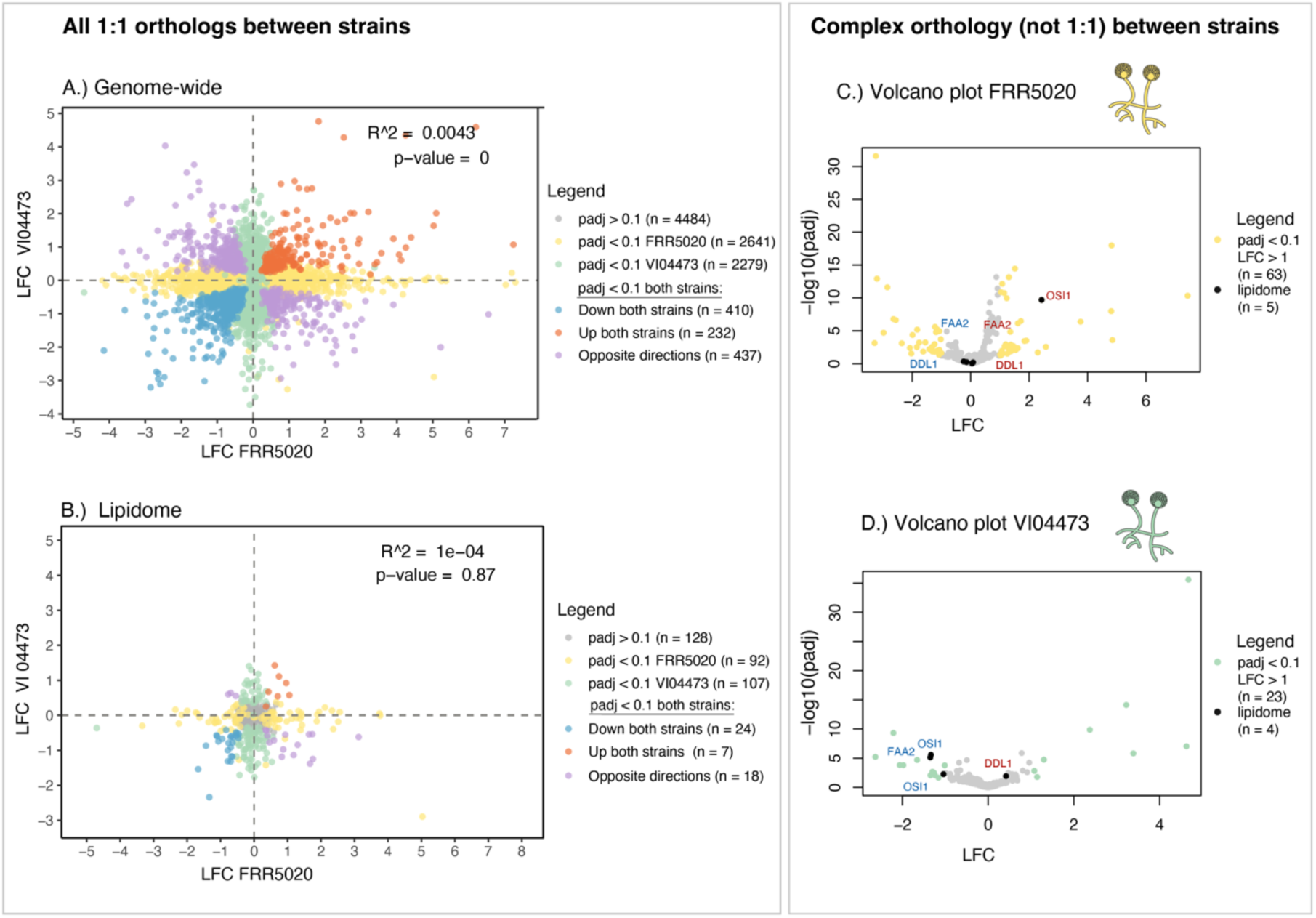
Gene expression results. Correlation of LFC between VI04473 and FRR5020 for all 1:1 orthologs in A.) and for lipidome orthologs in B.) Genes are grouped according to legend. LFC for genes with more complex, not 1:1 orthology relationships, is shown as volcano plots for C.) FRR5020 and D.) VI04473. Lipidome genes are highlighted in black.

Further, we wanted to investigate the regulatory differences between strains for genes specifically related to lipid and carotenoid synthesis. We annotated genes involved in lipid metabolism, i.e the ‘lipidome’ in FRR5020 and VI04473 based on orthology to 389 known lipidome genes in the strain MU402 (Sokołowska *et al*. 2023) (see methods). The lipidome includes genes related to carotenoid synthesis, these processes are therefore inextricably linked (Sokołowska *et al*. 2023). In concurrence with the genome-wide results, there was no correlation of LFC in the 371 lipidome 1:1 orthologs between strains (Figure 3B). Only 31 DEGs showed a similar direction of regulation, with 24 genes being upregulated and 7 being downregulated in both strains due to Ca starvation.

Due to strain-specific gene family expansion and contraction events, some genes are not in the set of 1:1 orthologs. We therefore investigated in more detail the genes with more complex gene orthology relationships (i.e not 1:1 orthologs between the two strains). 493 out of 601 such genes in FRR5020 and 371 out of 510 such genes in VI04473, respectively, have gene expression levels data to call DEGs. For these complex orthology genes there were 63 DEGs in FRR5020 (Figure 3C) and 23 DEGs in VI04473 (Figure 3D) with a LFC higher than 1 (padj < 0.1). Three lipidome orthogroups had significant DEGs in both strains, displaying different regulation of gene duplicates within and/or between strains; FAA2, DDL1 and OSI1 (Figure 3C-D; Supporting information). The orthogroup containing the FAA2 (medium chain fatty acid CoA ligase) gene family includes FRR5020 gene duplicates regulated in opposite directions and one downregulated VI04473 gene copy in response to Ca starvation. This could lead to less degradation of medium fatty acids. The second lipidome orthogroup includes the yeast ortholog DDL1 phospholipase, where the duplicates are also regulated in opposing directions in FRR5020, and a single gene in VI04473 is upregulated. DDL1 leads to the enhanced function of a vital protein synthesis protein, EF2 (Elongation factor 2), which is surprisingly downregulated in VI04473. The third orthogroup includes the yeast ortholog OSI1 (Oxidative stress induced 1), which is related to stress responses in yeast (Gruhlke et al. 2017). Both duplicates are downregulated in VI04473, whereas the single copy in FRR5020 is upregulated.

### Divergent regulation of lipid and carotenoids pathways under Ca starvation

There is a tradeoff in the production of some lipids (e.g. sterols, TAGs, fatty acids and phospholipids) and carotenoids as they derive from acetyl-CoA through the TAG, FAS and mevalonate pathways, respectively (Ranganathan *et al*. 2020). We therefore manually investigated the regulatory differences of these pathways under Ca starvation in the different strains (for a complete overview of candidate genes, see Supporting information). The mevalonate pathway leads to the synthesis of either carotenoids or lipids in the form of sterols (Avalos and Carmen Limón 2015; Sokołowska *et al*. 2023). Many of mevalonate genes were differentially expressed in one or both strains because of Ca starvation (Figure 4A). In the VI04473 the entire pathway is downregulated. There are three copies of hmgr, in which two are downregulated in VI04473. hmgr is a known rate-limiting step in the mevalonate pathway (Basson *et al*. 1986). In FRR5020 some of the genes upstream of FPP are downregulated, but not hmgr. After the formation of FPP there is a junction with clear divergent patterns between the two strains. In VI04473 the erg9 gene is downregulated, which could lead to less squalene, which is a precursor to various forms of sterols (Sokołowska *et al*. 2023). In FRR5020, one of the carG paralogs is downregulated leading to the carotenoid synthesis pathway. crgA is downregulated in FRR5020. crgA is a negative regulator of carotenogenesis by blocking transcription of carB and carRP. There are two additional copies of crgA in both strains, however they are not 1:1 orthologs. Neither of these paralogs are DEGs in either strain.

**Figure 4:**
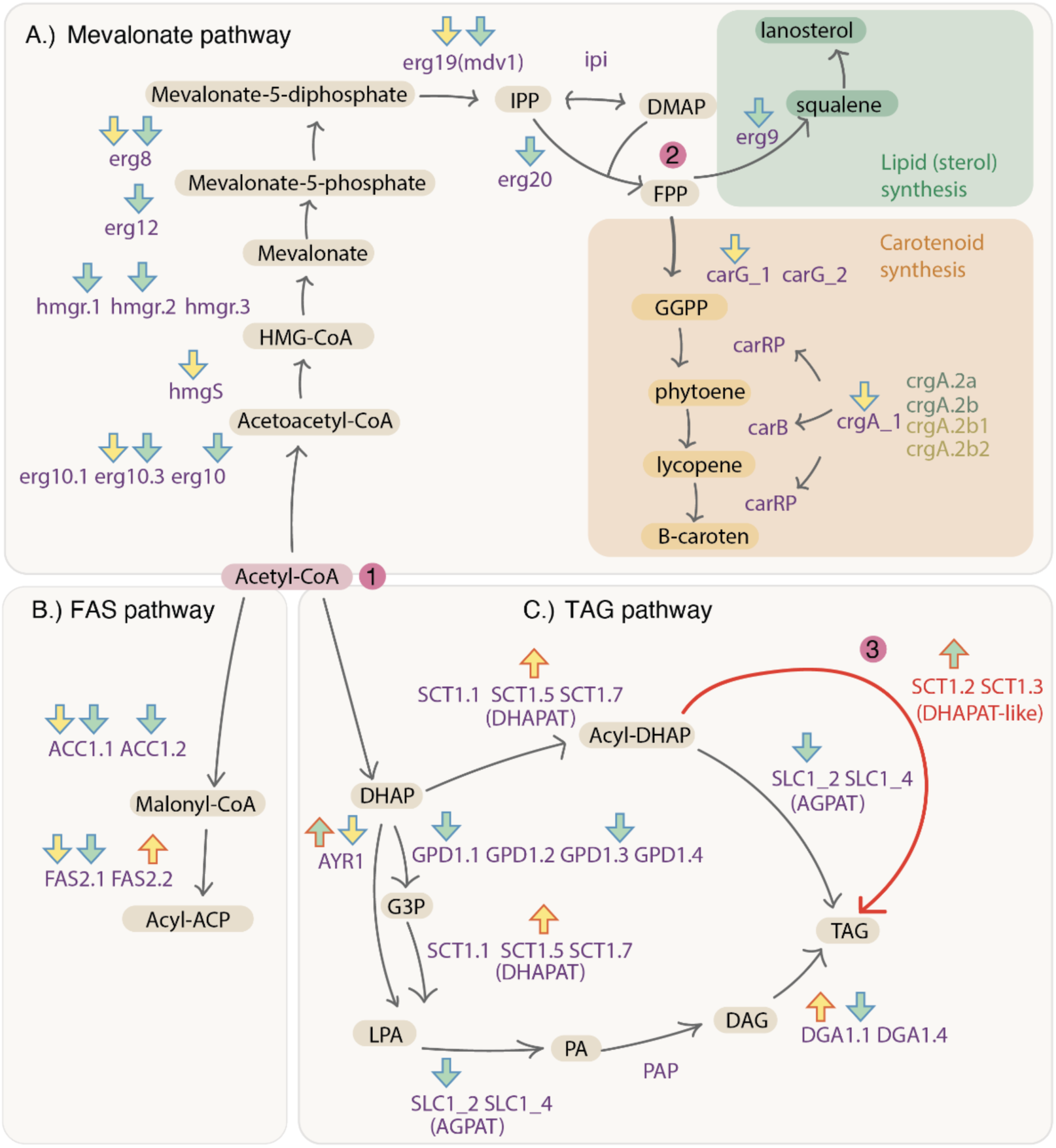
Key lipid pathway regulation under Ca starvation. A.) Mevalonate pathway B.) FAS pathway and C.) TAG pathway. Compounds are written in black, genes in purple. Arrows indicate if the gene is significantly down (blue outline) or upregulated (red outline), the fill of the arrow indicate strain, with FRR5020 in yellow and VI04473 in green. The numbers highlighted in pink illustrates key points: 1.) Acetyl-CoA can be utilized either in the mevalonate, FAS or TAG pathways. 2.) FPP can be utilized to produce sterols or carotenoids. 3.) indicates our proposed alternative pathway to TAG formation with the direct conversion of Acyl-DHAP via dhapat-like enzyme.

The downregulation of the mevalonate pathway in VI04473 is likely to free a lot of Acetyl-CoA for other lipid pathways. The FAS pathways is downregulated in VI04473, and mostly downregulated in FRR5020 with the exception of one of the FAS2 paralogs (Figure 4B). This could directly affect synthesis of TAGs and could lead to the accumulation of MAGs and DAGs (Figure 4B). Acetyl-CoA can also be used in the TAG pathway directly (Figure 4C). This pathway is a bit more complex to interpret. For VI04473 the downregulation of gdp1 and agpat would lead to less DAG and TAG. Interestingly, we detected five genes annotated as dhapat in both strains, with three being homologous to dhapat in *S. cerevisiae* (sct1.1, sct1.5 and sct1.7). However, two copies (sct1.2 and sct1.3) were in orthogroups with only Mucoromycota species and no *S. cerevisiae* ortholog – we call these dhapat-like. They contain a phospholipid/glycerol acyltransferase domain. In mammalian adipocytes, dhapat can lead to direct synthesis of TAG from Acyl-DHAP. We have indicated this route as a possible hypothesis for lipid accumulation in *M. circinelloides*. In the case of VI04473 this route would be a primary route for increase lipid accumulation under Ca deficiency.

### Regulatory differences between Ca-sensitive candidate genes

Ca is a ubiquitous signaling molecule which can have wide-ranging effects on gene regulation that is hard to disentangle (Roy *et al*. 2020). We therefore manually inspected candidate genes (Supporting information) to investigate genes known to be related to Ca levels. Several genes in the calcineurin signaling pathway were differentially regulated in both strains. A gene involved in calcium influx in response to environmental stresses, cch1 (Calcium Channel Homolog) (Locke *et al*. 2000), is upregulated in VI04473 and downregulated in FRR5020. Downstream of this gene is cmk21 (calmodulin-dependent protein kinase), which has four copies in both strains, with two being upregulated. One of the two calcineurin subunits, cnb, was downregulated in FRR5020. The crz1p transcription factor has putatively seven copies in both strains, and three of these are upregulated. Crz1p regulates a myriad of genes, including those involved in lipid metabolism (Cyert 2003). There are four copies of pmc1 (vacuolar Ca2+ ATPase), which is involved in depleting the cytosol of Ca in the presence of elevated Ca levels. These were, as expected, downregulated under Ca starvation in both strains. Similarly, pmr1 (plasma membrane ATPase related), which sequesters Ca into the Golgi apparatus, was downregulated in both strains under Ca starvation.

### No association between synteny and gene regulatory differences

The importance of gene order conservation in genome regulatory evolution has been the focus of much scientific debate (Poyatos and Hurst 2007; Kikuta *et al*. 2007; Bhutkar *et al*. 2008; von Grotthuss *et al*. 2010). We therefore investigated the role of genomic rearrangements on gene regulatory differences between FRR5020 and VI04473 using three different approaches. As above, we used both the Ca0 vs Ca1 and the P1 vs P4 contrasts to investigate gene expression differences between strains under more stressful vs less stressful conditions. First, we classified 1:1 orthologs between FRR5020 and VI04473 based on how many orthologous neighbors they share (either 0, 1 or 2 neighbors). If local microsynteny affects gene expression we would expect a higher correlation of LFC between strains for genes with 2 neighbors, than for 1 or 0 neighbors. We found that for the genes with 0, 1, or 2 there no differences in the correlation of LFC between strains, It should be noted that the sample size between the three groups is quite skewed (0 neighbors: n=160, 1 neighbor: n=2124, 2 neighbors: n= 6875). Second, we classified each gene based on the size of the synteny block it is in, and the distance to its closest synteny breakpoint. We expected that the ambiguous or strain-specific DEGs (Supporting information) would be located in less conserved and smaller synteny blocks or closer to a synteny breakpoints. However, we found no significant difference between the different DEG categories (Supporting information). Thirdly, we investigated only genes very close to synteny breakpoints, hypothesizing that the regulatory landscape of these genes is more likely to be affected by rearrangements. We investigated the difference in LFC between strains as a function of distance to the closest synteny breakpoint in genomic regions <10 000 bp from a breakpoint. We observed a weak, negative relationships between the two, but this is not significant. In conclusion, we find little evidence for genomic rearrangements between strains being mechanistically linked to the low level of gene regulatory conservation between the two strains.

## DISCUSSION

In line with two previous studies (Dzurendova *et al*. 2021; Dzurendová *et al*. 2021) we have demonstrated that Ca starvation leads to divergent phenotypic effects in the two *Mucor circinelloides* strains, with VI04473 responding to this stressor with increased lipid accumulation and FRR5020 responding with increased carotenoid production (Figure 1). We now also show that these strains have extremely divergent gene regulatory responses to Ca starvation (Figure 3), and there are extensive genomic rearrangements between the two strains (Figure 2).

In our study we find very low conservation of Ca-stress induced gene expression responses, both genome-wide and for lipidome genes (Figure 3) The lack of gene expression conservation could be specific to the type of stress elicited by Ca starvation. Given that we observed a much higher gene expression correlation between strains when we compared two conditions that represent low stress (P1 and P4, supporting information), this seems like a plausible explanation. This is concurrent with findings in yeast, where the conservation of gene expression under stress is dependent on the stressor, with heat shock responses being conserved whereas uracil limitation and rapamycin treatment had a higher number of genes with divergent expression (Guan *et al*. 2013). We found that the calcineurin signaling pathway was activated in both strains under Ca starvation. In both strains we found upregulation of multiple paralogs of the crz1 transcription factor under Ca starvation. This could trigger a cascade of responses affecting many downstream genes, including the dimorphic switch between yeast-like growth and filamentous growth (Lee *et al*. 2013), and could explain the lack of conservation of genome-wide expression across strains. However, the triggering of the calcineurin pathway by Ca depletion seem to be conserved over large evolutionary distances as a similar response has been shown in yeast to ensure survival of the cells under stress (Bonilla *et al*. 2002).

The overall conservation of expression of the lipidome is also not conserved between strains (Figure 3B). When we examined these large differences in expression in different pathways leading to the production of carotenoids and lipids, we found several plausible mechanisms explaining the phenotypic divergence between strains. Carotenoid production involves relatively few genes (Figure 4A), and we found that Ca starvation leads to the down regulation of a negative regulator of carotenogenesis, which could lead to higher production of carotenoids. Previous studies have documented that carotenoid production in *M. circinelloides* and *M. lusitanicus* is influenced by light (Naz *et al*. 2020), carbon source, temperature, salt concentration and oxygen level (Nagy *et al*. 2014). To the best of the authors’ knowledge, this study presents the first evidence of Ca deficiency as a novel trigger for carotenoid production through regulations of the mevalonate pathway in this organism. Moreover, this is the first study to report the co-regulation of carotenoid and TAG synthesis under Ca deficiency, providing groundbreaking insights for strain development and optimization aimed at the simultaneous production of lipids and carotenoids.

Disentangling lipid accumulation in VI04473 under Ca starvation is more complicated as there are multiple, interconnected metabolic pathways involved in lipid production. These pathways compete for Acetyl-CoA utilization, an important metabolic intermediate (Sokołowska *et al*. 2023). In VI04473 the mevalonate and FAS pathways are downregulated under Ca starvation, and it is therefore plausible that lipid accumulation is mainly due to TAG synthesis. However, the regulatory pattern of genes in the TAG network in VI04473 does not fit with the model of TAG synthesis in yeasts like *S. cervisiae*. We speculate whether TAG is formed directly from Acetyl-DHAP, (Figure 4C) . This pathway may represent an alternate route for lipid synthesis, more closely resembling lipid metabolism seen in mammalian adipocytes than in yeasts. Interestingly, similar mechanisms have been observed in algae, where Ca starvation also leads to increased lipid accumulation (Gorain et al., 2013). This indicates a potential common strategy across different groups of organisms for managing lipid storage under nutrient stress conditions, further highlighting the significance of Ca in regulating lipid metabolism. Whether or not this is homologous across these groups and lost in yeasts, or the product of recurrent evolution remains to be tested.

Synteny disruption between these two *M. circinelloides* strains seem to have no effect on the gene regulatory differences between strains, at least not due to Ca or P levels. Similar observations have been made in yeasts, where the co-expression of genes is a poor predictor of conservation of microsynteny or local gene neighborhoods (Poyatos and Hurst 2007). However, to get a better overview of the effect of genome rearrangements on gene regulation we would need to investigate gene expression differences over broader environmental conditions as well as include more species to reflect different evolutionary distances (Martin and Fraser 2018). Large-scale synteny comparisons have been carried out in other fungal lineages (Hane *et al*. 2011; Li *et al*. 2022), but never in Mucoromycota. Between FRR5020 and VI04473 most gene pairs have remained neighbors, we would therefore most likely not detect purifying selection to keep them linked to conserve co-expression (Poyatos and Hurst 2007). Furthermore, the extensive rearrangements we observe in *M. circinelloides* (Figure 3) could contribute to reproductive isolation and speciation in this genus, thus indirectly influencing divergence in evolutionary rewiring of gene regulatory networks. The extensive genomic rearrangements between VI04473 and FRR5020 is concurrent with comparisons between genomes of a high lipid-producing strain, WJ11, and a low lipid-producing strain, CBS 277.49, which also display large phenotypic differences (Tang *et al*. 2015b). We propose that it is possible that these strains may in fact represent different species. Investigating whether or not these two *M. circinelloides* strains mate and recombine, equivalent to what has been done for other strains (Wagner *et al*. 2020), would shed light on whether the rearrangements we observe (Figure 3) still segregate within *M. circinelloides* populations.

### Future perspectives

Our study highlights the value of genome-wide investigations of metabolic pathways, both to gain and deeper understanding of the evolution of the diverse *M. circinelloides* strains, but also to optimize production of valuable metabolites for industrial purposes. The value of such genomic studies have been well documented in the model species yeast *Saccharomyces cerevisiae* (Hittinger *et al*. 2015; Riley *et al*. 2016). For most species of Mucoromycota we still have very little knowledge about their genome biology, despite their ubiquitousness in nature, ecological significance, medical relevance and biotech potential. A major challenge now is to link genomic variation to organismal phenotypes through functional genomic studies in species which remain underutilized by industry and science. Such studies can ultimately lead to a detailed understanding of gene networks that is needed for optimization and engineering of bio-based production.

## Materials and methods

### Fungal strains and inoculum preparation

The selection of *Mucor circinelloides* strains was based on the results of the previous study performed by Dzurendova et al., 2021. The FRR5020 strain was originally obtained in the Food Fungal Culture Collection (FRR; North Ryde, Australia) and the VI04473 strain was obtained from the Veterinary Institute (VI, Ås Norway). The stock cultures were stored at - 80°C and consisted of asexual fungal spores in a glycerol-NaCl solution. To recover fungal culture from the cryo-vials and prepare spore inoculum, cultivation on malt extract agar was done. Malt extract agar (MEA) plates were prepared by dissolving 50 g/L of agar powder (VWR Chemicals - Agar Powder) in deionized water, which was then autoclaved at 115°C for 15 min and subsequently cooled down to 45°C before being poured into Petri dishes (VWR, 9 cm diameter). The MEA Petri dishes were stored at 4°C until use. Before cultivation, the MEA plates were exposed to UV light in a Class 2 biological safety cabinet (Cellegard ES) for 15 min in order to prevent any contamination. Three 5 µL droplets of stock spore suspension (stored in a -80°C freezer) were inoculated onto each MEA plate. The MEA plates were then incubated at 25°C for 4 days in a VWR Inco-line 68 R heat locker. The cultivation on MEA plates was done in four replicates to obtain enough spores for the experiments. For inoculum preparation, 10 mL of sterile saline solution was added to the MEA plates, then spores were harvested by mixing and scraping with single-use plastic loops, and then the spore solution was transferred into sterile Falcon tubes. The obtained spore solution was used as inoculum to inoculate media with different Ca and phosphorus concentrations.

### Cultivation under different Ca and phosphorus availability

To investigate the effect of Ca on pigment and lipid production in *Mucor circinelloides* strains FRR5020 and VI04473 under varying phosphorus availability, nitrogen-limited cultivation media were designed using a full factorial approach. The experimental design included three concentrations of inorganic phosphorus (Pi), provided as phosphate salts (KH₂PO₄ and Na₂HPO₄), and two Ca conditions: Ca1 (presence) and Ca0 (absence). The nitrogen-limited broth media were used as described by (Dzurendova *et al*. 2021), and contained the following components (g/L): glucose, 80; (NH₄)₂SO₄, 1.5; MgSO₄·7H₂O, 1.5; CaCl₂·2H₂O, 0.1; FeCl₃·6H₂O, 0.008; ZnSO₄·7H₂O, 0.001; CoSO₄·7H₂O, 0.0001; CuSO₄·5H₂O, 0.0001; MnSO₄·5H₂O, 0.0001. All chemicals were sourced from Merck (Germany). The reference phosphorus concentration (Pi1) was set at 7 g/L KH₂PO₄ and 2 g/L Na₂HPO₄, based on its frequent use in the cultivation of oleaginous *Mucoromycota* (Kosa *et al*. 2018). In addition to Pi1, media were formulated with a higher phosphorus level (Pi4, 4× Pi1) and a lower level (Pi0.5, 0.5× Pi1) to assess the impact of varying Pi availability. Cultivation was carried out using a microtiter plate system (Duetz-MTPS). The microtiter plate system (Duetz-MTPS) consists of deep well 24-square microtiter plates, sandwich covers, and clamp systems to mount microplates on a shaking platform of an incubator. In each well of the microtiter plates, 7 mL of media (Ca1P1, etc.) was inoculated with 5 μL of spore solution, and in total of 3 wells corresponding to biological replicates per treatment were prepared. The microtiter plates were mounted on the shaking platform using clamp systems in a shaker (Kuhnershaker X, Climo-shaker ISF1-X) and incubated at 25°C and 250 rpm for 5 days.

### Biomass preparation for analysis

After the cultivation in Duetz-MTPS, fungal biomass was washed with deionized water using vacuum filtration system. Washed biomass was transferred into sterile Eppendorf tubes (2 mL) and immediately placed on dry ice to prevent biomass degradation. All biomass samples were stored at -80°C until RNA extraction and chemical analysis. To freeze-dry the biomass for chemical analysis the tubes with frozen biomass were put into the dryer and maintained at 0.08 bar and -50°C for 2 days for the water to evaporate. The freeze-dried biomass was kept in the freezer at -20°C.

### Vibrational spectroscopy analysis

Before Fourier transform infrared (FTIR) spectroscopy analysis, the dried biomass was homogenized: 2 mL screw-cap microcentrifuge tube was filled with 250 mg (710-1180 um diameter) acidwashed glass beads (Sigma Aldrich, USA) and a small amount of biomass was then added to the tube, followed by 0.5 mL of distilled water. The tubes were placed in a Precellys Evolution tissue homogenizer (Berlin Instruments, France) at 5500 rpm, 20 sec cycle length, and 6 cycles (2x20 s, 3 runs) to ensure complete homogenization of the biomass. Approximately 10 μL of homogenized biomass was pipetted onto an IR transparent 384-well silica microplate, with one free spot between each sample. Three technical replicates were pipetted onto 384-well FTIR silica microplate, for each sample. After drying at room temperature for 30-45 min the samples were measured in FTIR spectrometer. FTIR spectra were recorded in transmission mode using a high-throughput screening extension (HTS-XT) unit coupled to a Vertex 70 FTIR spectrometer (both Bruker Optik GmbH, Leipzig, Germany). Spectra were recorded in the region between 4000 and 500 cm−1, with a spectral resolution of 6 cm−1, a digital spacing of 1.928 cm−1, and an aperture of 5 mm. For each spectrum, 64 scans were averaged. Spectra were recorded as the ratio of the sample spectrum to the spectrum of the empty IR transparent microplate. In total, 63 FTIR biomass spectra were obtained. The OPUS software (Bruker Optik GmbH, Leipzig, Germany) was used for data acquisition and instrument control.

FTIR spectra were analyzed using Orange 3.16 (Demšar *et al*. 2013). For the estimation of lipid to protein ratios, FTIR spectra were preprocessed using extended multiplicative signal correction (EMSC) with linear, quadratic, and cubic terms (Forfang *et al*. 2017). The calculations of lipid-to-protein ratios were done in Microsoft Excel (Microsoft Excel 2019, Microsoft Corp., Redmond, WA, USA) by dividing the relative absorbance values for the -C=O stretching peak at 1,745 cm^−1^ (correlate with the total lipid content in cells) by the Amide 1 peak at 1,650 cm^−1^ (reference band for biomass).

In order to determination of the total relative carotenoid content in the biomass, analysis using a FT-Raman spectroscopy was performed. The freeze-dried biomass was transferred into 2 mL screw vials with 400 μL glass inserts until the bottom of the vials was completely covered with biomass, making three technical replicates for each sample. The samples in glass vials were put into a 96-well sample box, for 400 μL flat bottom glass insert and then placed onto a V-grooved sample mount with the samples facing the lens assembly in the MultiRAM FT-Raman spectrometer (Optic GmbH, Germany). The first and the last sample vials contained acetonitrile, used for reference measurements.

FT-Raman spectra were analyzed using Orange 3.16 (University of Ljubljana, Slovenia) (Demšar et al., 2013, Toplak et al., 2017). For the estimation of the relative carotenoids content, FT-Raman spectra were preprocessed using extended multiplicative signal correction (EMSC) with linear, quadratic, and cubic terms (Dzurendová *et al*. 2021). The calculations were done in Microsoft Excel (Microsoft Excel 2019, Microsoft Corp., Redmond, WA, USA) by dividing the Raman intensities at 1,523 cm^-1^ (related to carotenoids) and 1,445 cm^-1^ (related to biomass).

### Lipid extraction and fatty acid profile analysis

Analysis of the total lipid content and fatty acid profile was performed according to methodology in (Langseter *et al*. 2021). 20 mg of the freeze-dried biomass together with approximately 250 mg (710-1180 um diameter) of acid-washed glass beads (Sigma Aldrich, USA) were filled in a 2 mL screw-cap polypropylene tube. 500 μL chloroform was added to the mix, and 100 μL of the internal standard was pipetted with a Hamilton syringe. The mix was then homogenized in a Precellys Evolution tissue homogenizer (Berlin Instruments, France) at 5500 rpm, 20 s cycle length, and 6 cycles (2x20 s, 3 runs). The biomass was transferred into glass reaction tubes by washing the polypropylene tube 3 times with 800 μL methanol-chloroform-hydrochloric acid solvent mixture (7.6:1:1v/v). 500 μL of methanol was added into the glass reaction tubes after washing. The mixture was incubated at 90°C for 90 min in a Stuart SBH130D/3 block heater (Cole-Parmer, UK). After the samples were cooled down to room temperature, a small amount of sodium sulphate was added together with 1 mL of distilled water. The mixture was cooled down to room temperature after incubation. To extract the fatty acid methyl esters (FAMEs), 2 mL of hexane was added, and vortex mixed for 10 s before centrifugation at 3000 rpm for 5 minutes at 4°C. The upper organic phase was collected in glass tubes. The upper phase was extracted two more times, in addition, a 2 mL hexane-chloroform mixture (4:1 v/v) was added. To evaporate the solvent, the glass tubes were placed in a Stuart SBH130D/3 block heater (Cole-Parmer, UK) connected to nitrogen at 30°C. The FAMEs were transferred into GC vials by washing the glass tubes 2 times with 750 μL hexane (containing 0,01% butylated hydroxytoluene (BHT, Sigma-Aldrich, USA)), followed by pipette mixing approximately 15 times. The GC vials were stored in a -20°C freezer until GC-FID analysis.

Determination of total lipid content and fatty acid composition was performed by using gas chromatography 7820A System (Agilent Technologies, USA), equipped with an Agilent J&W 121–2323DB-23 column, 20m × 180 μm × 0.20 μm and a flame ionization detector (FID). Helium was used as a carrier gas. The total run time for one sample was 36 min with the following oven temperature increase: initial temperature 70 °C for 2 min, after 8 min to 150 °C with no hold time, 230 °C in 16 min with 5 min hold time, and 245 °C in 1 min with 4 min hold time. The injector temperature was 250 °C and 1 μL of a sample was injected (30:1 split ratio, with split flow 30 mL/min). For the identification and quantification of fatty acids, the Supelco 37 Component FAME Mix (C4–C24 FAME mixture, Sigma-Aldrich, USA) was used as an external standard, in addition to C13:0 TAG internal standard. Measurements were controlled by the AgilentOpenLAB software (Agilent Technologies, USA).

### DNA isolation and whole genome sequencing

DNA isolation started from flash-frozen fungal tissue. Sample was homogenized by grinding under liquid nitrogen. DNA was isolated using the Nucleobond HMW DNA extraction kit (Macherey-Nagel) following the manufacture protocol with small modification. For lysis sample was incubated at 50°C for 2.5 hours and all centrifugation step was done at minimum 10 000g. Quality of the DNA was evaluated by running the sample on 0.5% agarose gel and by using Nanodrop 8000 Spectrophotometer. After DNA isolation, size selection was performed by using SRE kit (PacBio). ONT library preparation was done using the Native Barcoding Kit 24 V24 (SQK-NBD114.24) (Oxford Nanopore) following the manufacture protocol. The library was loaded on an R9.4.1 flow cell and was sequenced using a Promethion24 device. To maximize the data output, we performed a nuclease flush when the ratio of the sequencing pores dropped under 20%. Nuclease flush and library reloading were repeated twice. Guppy 5.1.13 was used for base calling with the High-accuracy base calling model.

### Genome assembly and annotation

The genome assembly was done by first by removing short reads (<4000bp) and low-quality ones (q<7), then assembled with Flye version 2.9.1 (Kolmogorov *et al*. 2019) and Canu version 2.2 (Koren *et al*. 2017). The default correctedErrorRate for Nanopore reads is 0.144, which might be too conservative for the newer chemistries of ONT. We therefore tested out several values for this parameter (default =0.144, 0.1, 0.05 and 0.03). The assemblies were then evaluated with Inspector version v1.0.2 (Chen *et al*. 2021) and Merqury version v1.3 (Rhie *et al*. 2020) for accessing the quality (QV) and completeness; and with BUSCO version 5.4.3 (Manni *et al*. 2021) for accessing the quantity of conserved genes. Based on assessing correctness, contiguity, and completeness for all assemblies we selected the Canu assemblies with correctedErrorRate=0.03 (supporting information).

Annotation was done with the funannotate pipeline version v.1.8.13, (Palmer and Stajich, 2022) using the RNA-seq data generated in this project. This pipeline was developed to annotate fungi and is widely used. RNA-seq samples were merged to the corresponding strain to serve as evidence for the annotation. The assemblies were first cleaned to remove the repetitive contigs, then masked the repeat regions using the default masker tantan in funannotate. The gene models were predicted with several methods and then combined into one by Evidence Modeler. Functional annotation was run with all the implemented tools in the pipeline: Pfam, InterProScan, Eggnog, UniProtKb, Pobius, antiSMASH, MEROPS. Centromeres were manually annotated using BLAST+ v.2.14.1 (Altschul *et al*. 1990) with sequences of known centromere motifs and centromere-specific transposable elements (GREM-LINEs) from (Navarro-Mendoza *et al*. 2019).

### Comparative Genomics

To investigate the phylogenetic placement of our two *Mucor circinelloides* strains as well as gene family evolution we used Orthofinder2 (Emms and Kelly 2015, 2019), based on protein sequence data from all genes in a set of species. We downloaded protein sequence data based on genome annotations from the Ensemble collection number 54 for Mucoromycota (http://ftp.ensemblgenomes.org/pub/fungi/release-54/fasta/fungi_mucoromycota1_collection/). This included an additional strain of *M. circinelloides* we denoted ad MucCirc4. We also downloaded the protein sequences from *M. circinelloides f. lusitanicus* CBS277.49 (Navarro-Mendoza *et al*. 2019), hereafter denoted as MU402. We excluded *Mortierella verticillata* (GCA000739165) because it contained very few genes (1523). As outgroup species we included *Conidiobolus coronatus*, *Aspergillus nidulans* and *Saccharomyces cerevisiae.* Including FRR5020 and VI04473 this resulted in a dataset of a total of 30 species (see supporting information for assembly versions and data location).

We plotted synteny between *M. circinelloides* strains for which we had chromosome-level assemblies; FRR5020, VI04473 and MU402 using GENESPACE v.1.4 (Lovell *et al*. 2022). A synteny riparian plot was generated using VI04473 as a reference. In addition, pairwise dot plots were generated between all three assemblies (supporting information).

### RNA isolation and transcriptomic sequencing

RNA was isolated for both strains and both experiments (6 + 2 treatments, 4 replicates for each). The RNA extraction was performed with the RNeasy Plus Mini kit. Concentration and integrity of RNA were determined using the 4150 TapeStation system and Nanodrop 8000 (Thermo Fisher Scientific). Most samples had RIN values higher than 6. Preparation of sequencing libraries and RNA sequencing was carried out at Novogene. Illumina sequencing libraries (150 PE) were prepared with the TruSeq Stranded mRNA Library kit according to the standard protocol. Libraries were sequenced using the Illumina Hiseq 2500 platform.

### Differential gene expression analyses

We obtained sufficient RNA-seq data for at least 3 replicates from each treatment from each strain (Supporting information). RNA sequencing data were processed for each strain separately using the nfcore/rnaseq v3.11.2 pipeline (Ewels *et al*. 2020). This pipeline includes the following steps: Raw reads underwent quality and adapter trimming with Trim Galore, removal of genome contaminants with BBSplit, removal of ribosomal RNA with SortMeRNA, alignment to their respective genome assembly using STAR and quantification with Salmon. Alignments were sorted and indexed with SAMtools, UMI-based deduplication was carried out with UMU-tools, and duplicates were marked with picard MarkDuplicates. StringTie was used for transcript assembly and quantification and bigwig coverage files were generated with BEDTools and bedGraphToBigWig. Extensive quality control was carried out for each sample and collated with MultiQC. Differential expression analyses were carried out using DESeq2 package (Love *et al*. 2014) in R. We compared gene expression between strains using 1:1 orthologs defined by Orthofinder.

### Candidate gene analysis

We used lipidome genes annotations for the strain MU402 in (Sokołowska *et al*. 2023). To annotate lipidome genes in FRR5020 and VI04473 we identified orthogroups that contained MU402 lipidome genes. FRR5020 and VI04473 genes within those orthogroups were assigned as lipidome orthologs of the closest MU402 lipidome gene based on the orthogroup tree distance. We manually inspected candidate genes known to be related to Ca levels, lipids and carotenoid pathways (Supporting information). Some of the genes were not properly annotated in both strains – these were manually annotated using BLAST.

## Author contributions

The study was conceived and planned by SRS, VS and HTB. Experiment was carried out by VB, IEL and DB. DNA isolation and sequencing was performed by MA, RNA isolation by VB, IEL and JF. THT did genome assembly and annotation. HTB did comparative genomics analyses and differential expression analyses. HTB and LG carried out synteny analyses. Analyses of regulatory networks was done by HTB and VS. SRS, VS and HTB wrote this manuscript with help and contributions from all authors.

## Acknowledgements

All laboratory work was carried out at the Centre for Integrative Genetics (CIGENE) laboratory and REALTEK laboratory at the Norwegian University of Life Sciences (NMBU). The computations were performed on resources provided by Orion High Performance Computing Center (OHPCC) at NMBU as well as Sigma2 - the National Infrastructure for High-Performance Computing and Data Storage in Norway. We are grateful to Anna Muszewska for providing with lipidome annotations.

