## Supporting information for "Calcium starvation leads to strain-specific gene regulation of lipid and carotenoid production in *Mucor Circinelloides*"

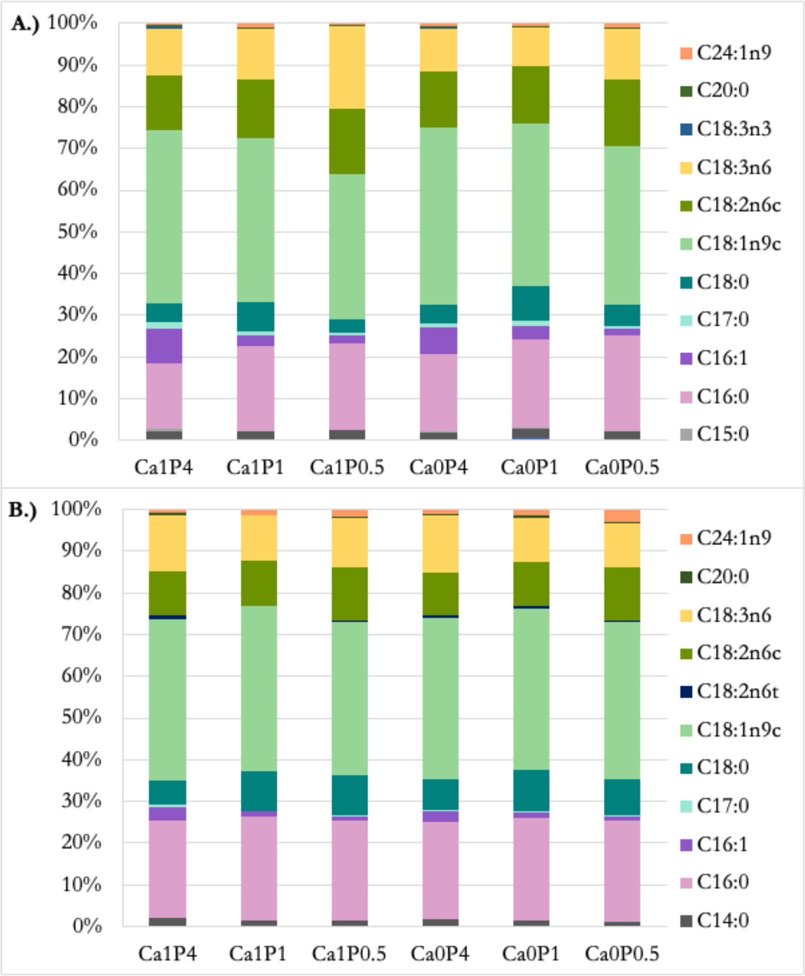


Figure S1: Fatty acids profiles for A.) VI04473 and B.) FRR5020 at the six treatments with different combinations of Ca and P levels. Each fatty acid is colored according to legend.

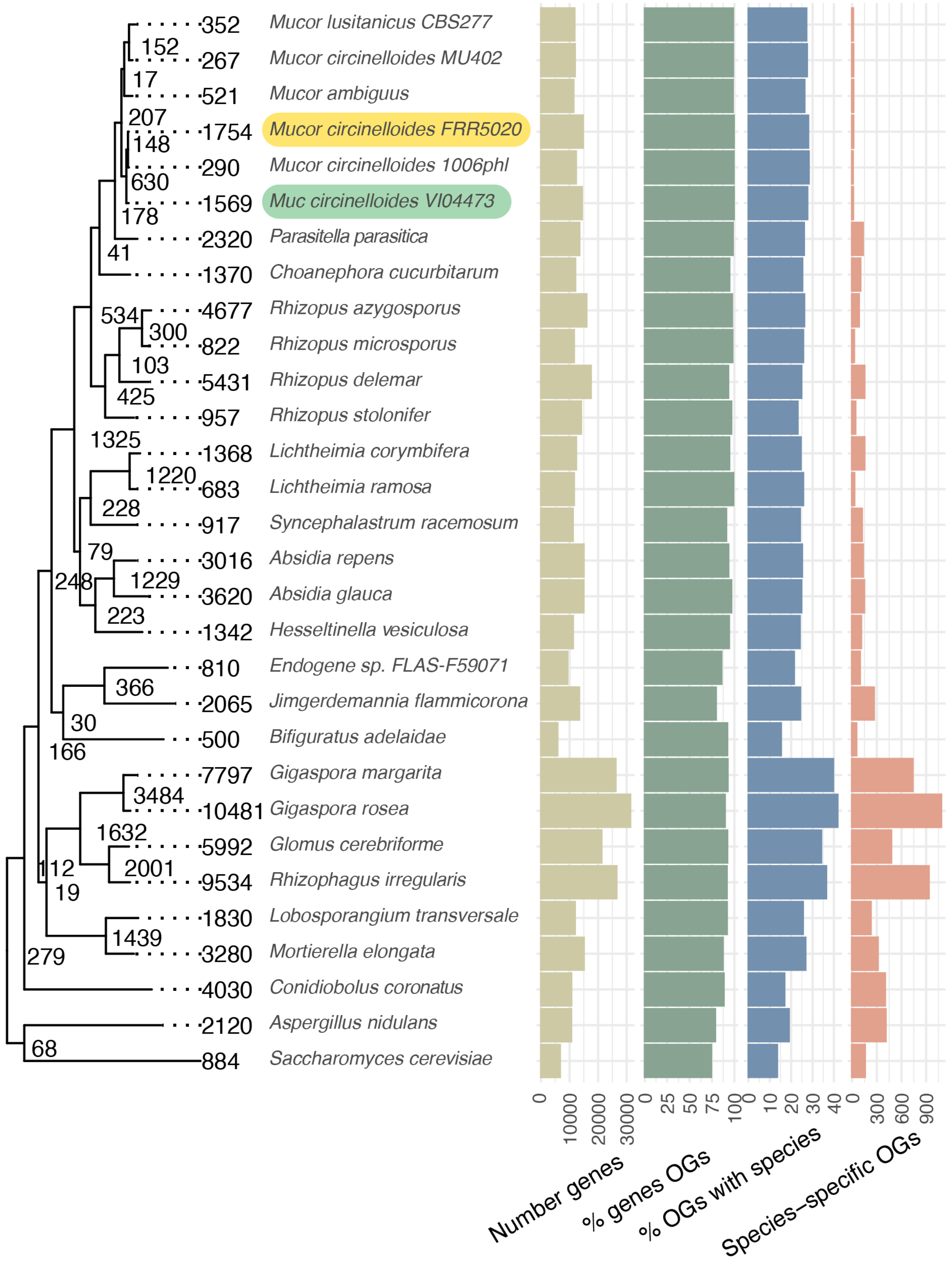


**Figure S2: Phylogenetic tree and Orthofinder results for 30 strains/species of fungi Mucoromycota.** The mains strains investigated in this study highlighted in yellow (FRR5020) and turqoise (VI04473). Phylogenetic tree was generated by Orthofinder. The number of gene duplications is denoted at each tip and node. Barplots show the number of genes placed in orthogroups (OGs), the percentage of genes in orthogroups, the percentage of orthogroups containing that species, and how many species-specific orthogroups there are.

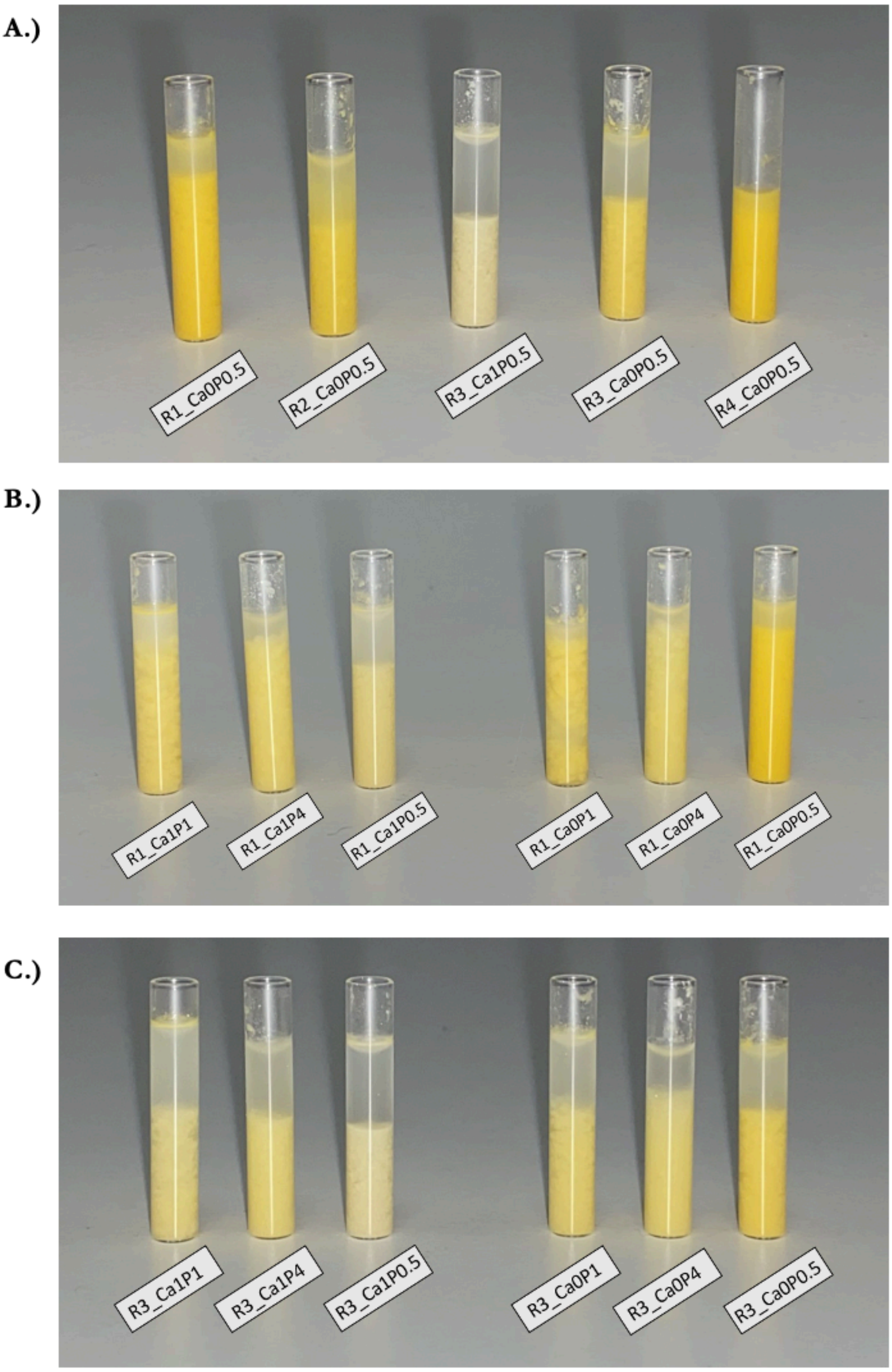


**Figure S3:** Pictures of *Mucor circinelloides* strain FRR5020 tissue samples at different combinations of Ca and P treatments. R denotes replicate.

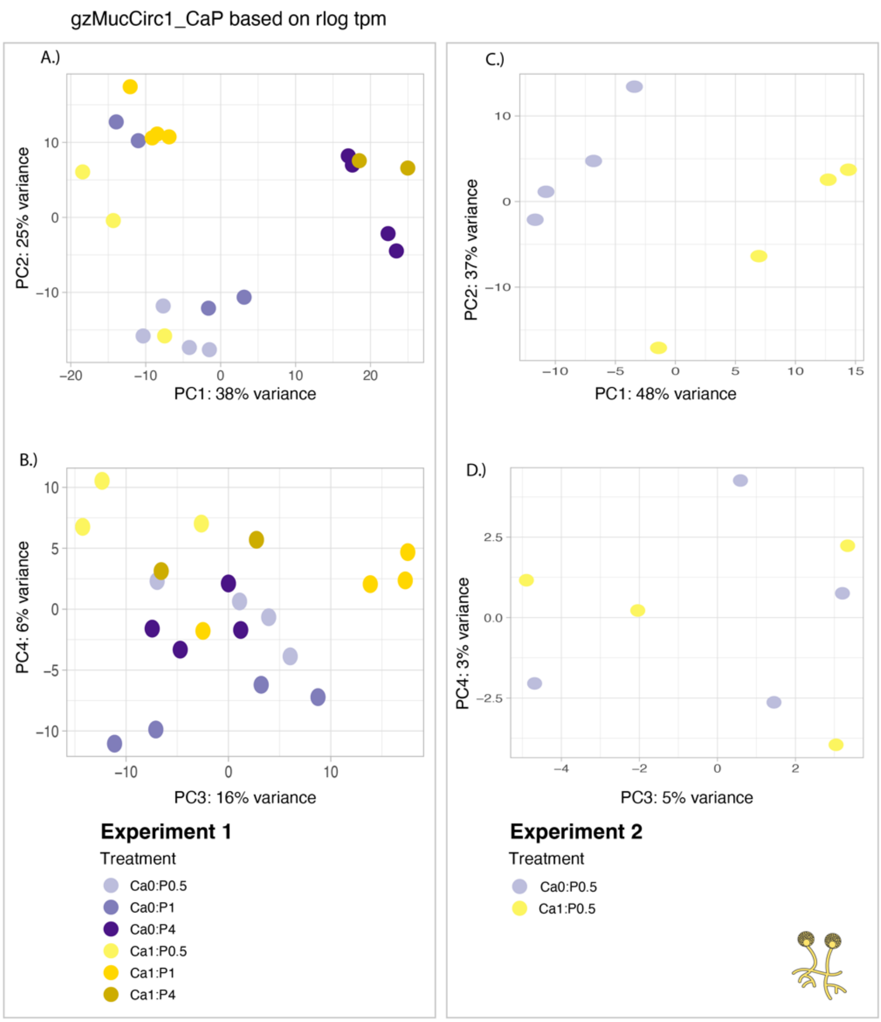


**Figure S4:** PCA plots for FRR5020 from experiment 1 (Ca and P) and experiment 2 (only Ca). Samples are colored according to legend.

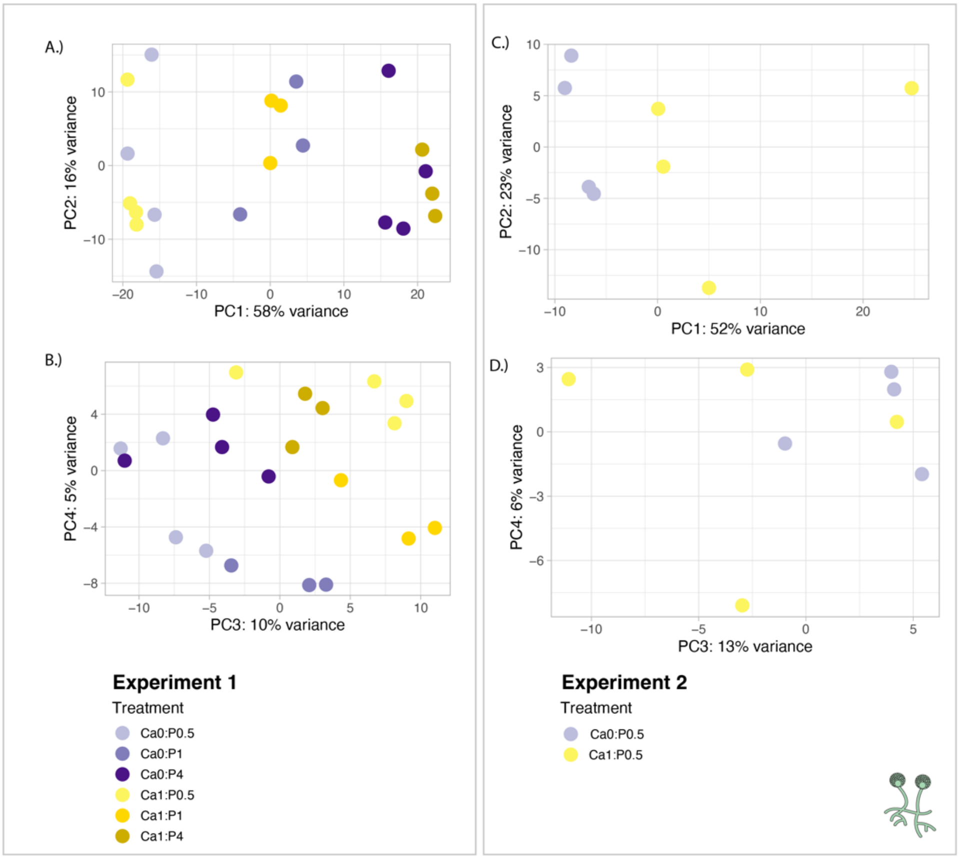


**Figure S5:** PCA plots for VI04473 from experiment 1 (Ca and P) and experiment 2 (only Ca). Samples are colored according to legend.

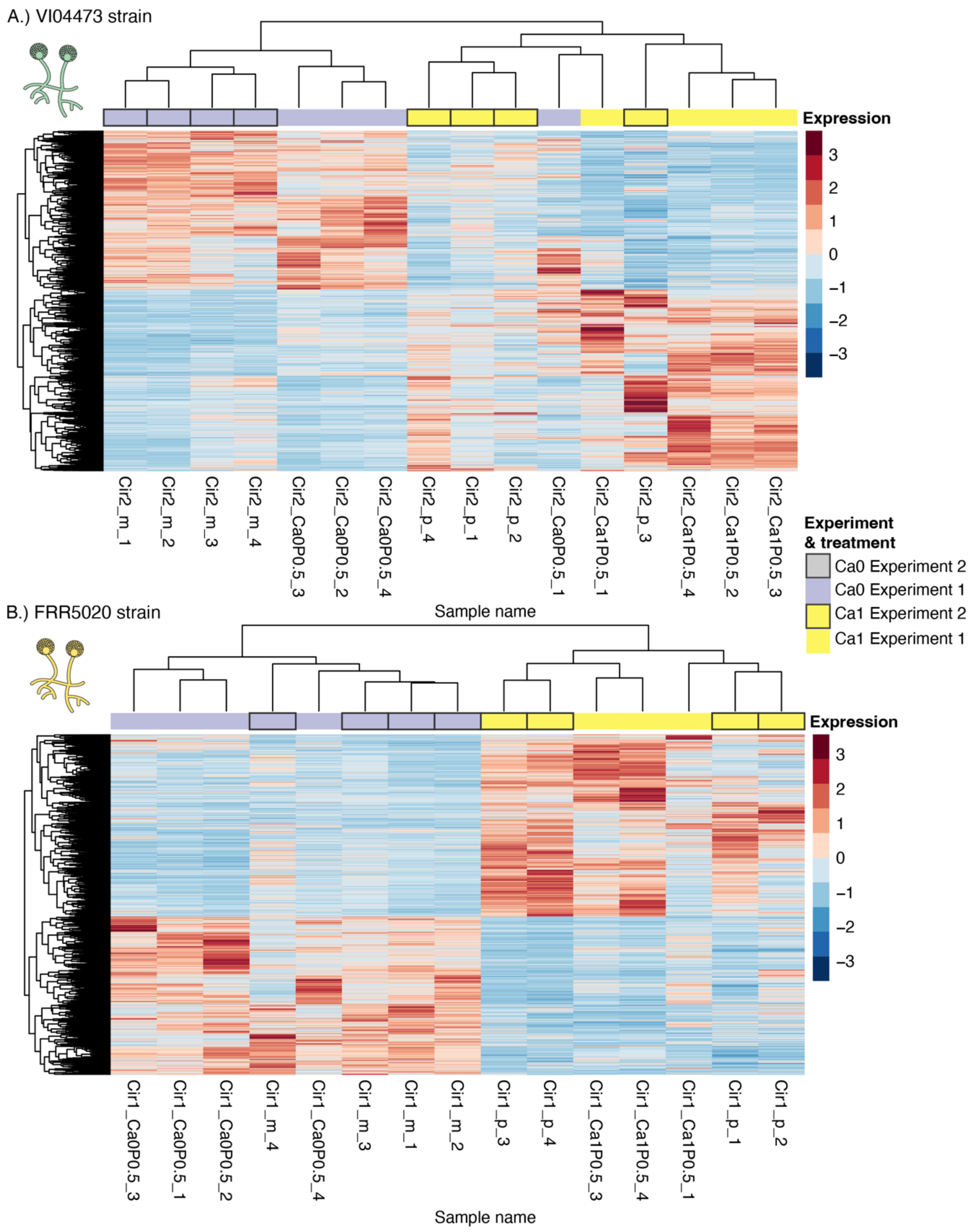


**Figure S6:** Heatmaps of gene expression of significant DEGs (padj < 0.1) in *Mucor circinelloides* strains A.) VI04473 and B.) FRR5020. Differential expression (log2FoldChange) is colored according to legend. Samples are from two separate experiments; Experiment 1 and Experiment 2, with two treatments; Ca0 or Ca1, according to legend. All samples have P0.05. Sample names for the different technical replicates are shown at the bottom.

Table S1: Overview of ortholog relationships between VI04473 and FRR5020 based on Orthofinder results.

| Ortholog relationship | Number of genes |
|---|---|
| **1:1** | 10430 |
| **1 FRR5020 : many VI04473** | 133 |
| **many FRR5020 : many VI04473** | 53 |
| **1 VI04473 : many FRR5020** | 248 |
